# Visual speech supports phonetic attunement in the absence of early auditory input

**DOI:** 10.64898/2026.05.26.727650

**Authors:** Alessandra Federici, Marta Fantoni, Chiara Battaglini, Francesca Collesei, Eva Orzan, Janet F. Werker, Linda Polka, Benedetta Bianchi, Giacomo Handjaras, Giovanni M. Di Liberto, Davide Bottari

**Affiliations:** MoMiLab, IMT School for Advanced Studies - Lucca, Italy; Neurolinguistics and Experimental Pragmatics (NEP) Lab, University School for Advanced Studies IUSS Pavia, Italy; IRCCS Materno Infantile Burlo Garofolo, Trieste, Italy; Department of Psychology at the University of British Columbia, Vancouver, Canada; Faculty of Medicine and Health Sciences, School of Communication Sciences and Disorders, McGill University, Montreal, Canada; IRCCS Meyer, Azienda Ospedaliero-Universitaria Meyer, Firenze, Italy; School of Computer Science and Statistics, University of Dublin, Trinity College, Ireland; ADAPT Centre; Trinity College Institute of Neuroscience, University of Dublin, Trinity College, Dublin 2, Ireland; Trinity Centre for Biomedical Engineering, University of Dublin, Trinity College, Dublin 2, Ireland

## Abstract

The first year of life is considered a sensitive period for the acquisition of phonetic categories, a hallmark of successful native language specialization. The extent to which this process depends on early auditory experience and intrinsic biological constraints remains unresolved. We measured neural encoding of continuous natural speech in hearing children (HC) and cochlear implant (CI) users with congenital or acquired deafness, contrasting children with and without access to auditory input in the first year of life. Speech encoding was present across all groups, but its specificity depended on early input: auditory phonetic features were encoded only in children exposed to speech within the first year, whereas visually discriminable phonetic features were encoded regardless of auditory deprivation. These findings show that early sensory input gates phonetic attunement; this constraint is not limited to or grounded in audition but instead reveals a sensitive period that is modality-flexible in mechanism and experience-dependent in expression.

## Introduction

The anatomical structures that enable speech are universal, and all languages have evolved around shared core articulatory mechanisms that the brain can interpret^1,2^. Yet, despite this shared biological foundation, speech perception becomes rapidly specialized for the native language during infancy. Early sensory experience is known to tune the brain to the acoustic structure of the native tongue, including its phoneme inventory ^3,4^. Until approximately 10 months of age, infants remain sensitive to non-native speech sound contrasts, discriminating minimal pairs that distinguish meaning in languages other than their ambient linguistic environment ^5^. With experience, perception becomes specialized for the language(s) they are exposed, refining discrimination of native phonetic patterns while losing sensitivity to non-native contrasts ^6^. This early developmental shift is widely attributed to the statistical distribution of speech sounds during the first year of life ^7^. Depending on the characteristics of the native language, distinct phonetic categories may or may not emerge. A classic example is the /r/-/l/ contrast English (e.g., rock vs. lock), which is absent in Japanese. This contrast is discriminated by young but not older Japanese infants in the first year of life and remains challenging for Japanese adults ^8–10^.

The shaping of the neural circuits responsible for phonetic contrast processing occurs in response to experience during a sensitive period of development ^11^. Brief auditory exposure to different phoneme distributions within this plastic window strongly shapes phonetic contrast processing, whereas after this period native phoneme categories become more resistant to change ^12^. Converging evidence indicates that the timing of this sensitive period is regulated by biological constraints. When brain maturation is pharmacologically advanced, as in the case of infants exposed in utero to serotonin reuptake inhibitors, which exert GABAergic effects, an earlier onset of the sensitive period for native phonetic categories is observed ^13^, indicating that the timing of this maturational shift is malleable. By contrast, preterm infants do not exhibit earlier development of categorical phonetic perception ^14^, suggesting that increased speech exposure alone does not accelerate its onset. However, experience can also modulate this timeline: on the one hand, exposure to more complex or variable input, as experienced by bilingual infants, appears to extend the sensitive period ^15^; on the other hand, infants of untreated depressed mothers, who may provide reduced linguistic input, show evidence of a delayed opening of the sensitive phase for phonetic perceptual attunement ^13^. Together, these findings indicate that both enriched and impoverished input can prolong its duration or postpone its emergence.

Although early sensory experience can modulate the timing and outcome of this sensitive period, a key unresolved question is whether specific sensory channels provide the determinant input for phonetic attunement. In particular, it remains unclear whether auditory input gates the sensitive period for phonetic attunement, or whether other modalities could support the extraction of speech statistics during early development in case hearing is not available. Audition plays a more dominant role in speech processing than vision throughout the lifespan, and phonology is primarily sound-dependent ^1^. However, speech perception is inherently multimodal, and speech patterns are extracted through vision as early as two months of age ^16,17^. During the second half of the first year of life, infants increasingly attend to the mouth ^18^, and neural measures of continuous speech processing show enhanced speech encoding in the presence of visual information from a talking face ^19^. Importantly, discrimination of visual speech cues is also shaped by early linguistic experience during the first year of life ^20–22^, suggesting that the sensitive period for speech attunement extends beyond the auditory modality.

To investigate the role of sensory experience in shaping the sensitive period for phonetic perceptual attunement, we studied individuals with delayed access to auditory input. Infants with congenital bilateral profound deafness are auditorily deprived until they receive cochlear implants (CIs), which typically occurs in at the end of the first year of life. As a result, they are exposed to speech sounds only after the age at which the sensitive period for native phonetic attunement has typically closed in hearing children. Prior to implantation, however, they can access speech patterns through vision. Importantly, while some phonemic contrasts are readily distinguished via audition but difficult to distinguish visually (for instance, voicing contrast such as /b/ vs. /p/), others differ more subtly in sound, yet are more easily discriminable visually (for instance, place of articulation contrasts such as /m/ vs. /n/; ^23^). Contrasts with this trade-off in their auditory and visual salience are common across human languages ^24,25^. Leveraging this naturally occurring variation in auditory and visual speech information, we tested the hypothesis that the narrowing of the sensitive period of native phonetic perceptual attunement within the first year of life involves multiple sensory channels; it is not limited to or controlled by auditory experience alone. Accordingly, we predicted that congenitally deaf children whose auditory function was restored after this period would successfully encode phonemic contrasts supported by visual phonetic cues, but not those relying solely on acoustic phonetic cues. The alternative hypothesis posits audition gates the sensitive period for attunement to speech. This view predicts that phonetic attunement would occur later following cochlear implantation, once children gain access to auditory speech statistics.

We tested these ideas using a model-driven naturalistic neuroscience approach ^26^. We recorded electroencephalography (EEG) during continuous auditory and audiovisual speech in 93 children (the final sample comprised 75 of them), including a sample whose hearing was restored with CIs and age-matched hearing controls (HC). CI participants were classified according to the onset of bilateral profound deafness as congenitally deaf (CD) or as having acquired deafness (AD). Only CD children experienced early auditory deprivation, whereas AD individuals were exposed to auditory speech statistics during the typical sensitive period for phonetic perceptual attunement (see Figure 1A). This contrast allowed us to isolate the selective role of early auditory input during ontogeny.

**Figure 1.**
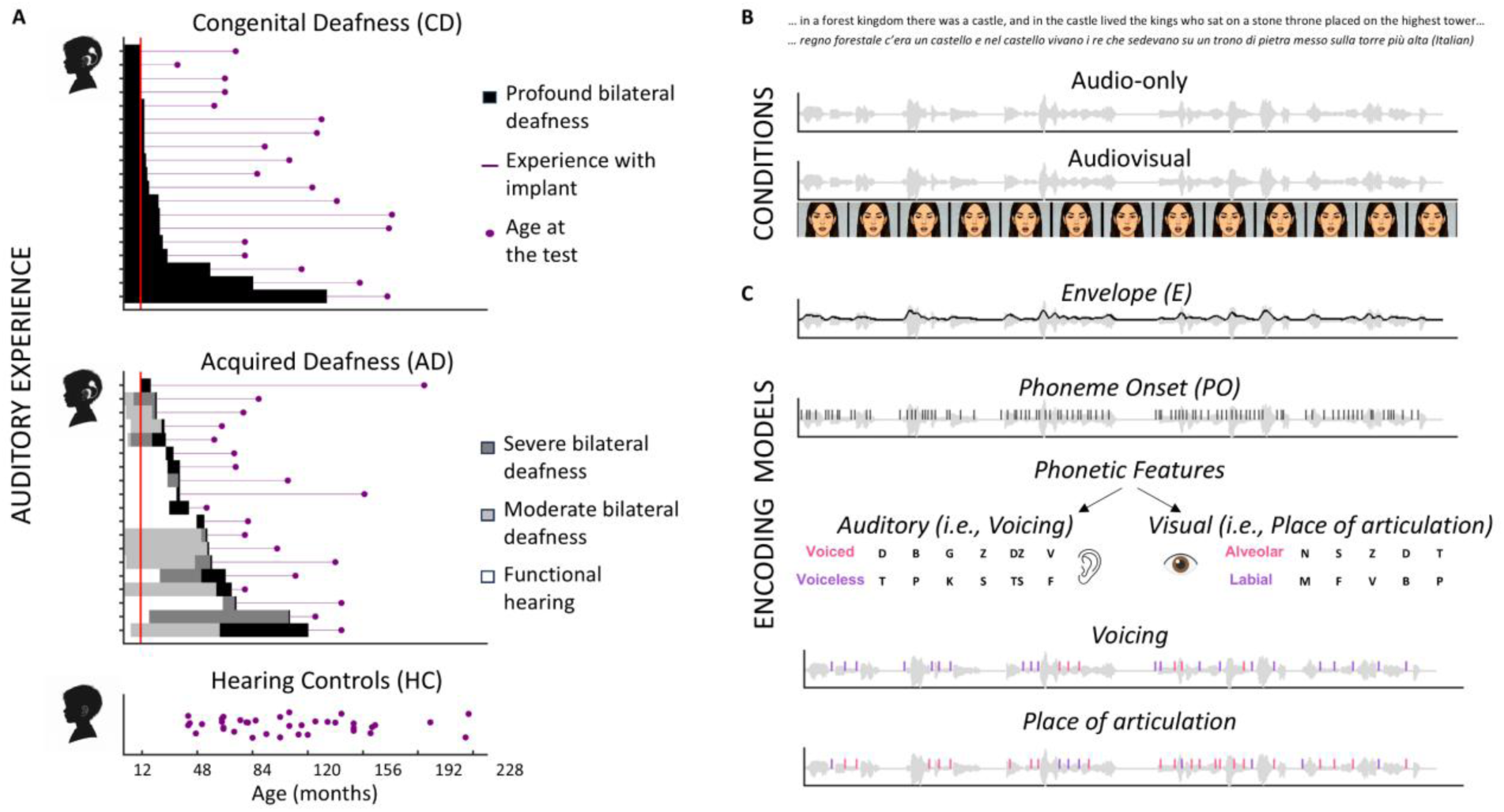
Auditory experience, experimental conditions, and encoding models. A) Auditory experience. The plots depict the auditory history of each participant up to the time of testing (magenta dots). For cochlear implant (CI) users, periods of hearing status are color-coded: black indicates profound bilateral deafness, dark grey severe bilateral deafness, light grey moderate bilateral deafness, and white functional hearing. Magenta lines represent the duration of cochlear implant use. The red line marks the end of the first year of life, highlighting the critical group difference: only children with congenital deafness (CD) experienced auditory deprivation during the first year, whereas children with acquired deafness (AD) had some residual functional hearing and were exposed to auditory input during early development. Age-matched hearing controls (HC) are shown at the bottom. B) Experimental conditions. Participants listened to a continuous narrative presented in two conditions: audio-only and audiovisual. In the audiovisual condition, the acoustic signal was accompanied by the speaker’s facial articulations. C) Encoding models. Predictors were progressively entered into multivariate encoding models to predict EEG activity. The baseline model included the speech envelope (E). A second model additionally included phoneme onset (PO). Finally, one phonetic feature was alternatively added: voicing (auditory feature) or place of articulation (visually salient feature).

Using multivariate encoding models, we predicted individual neural activity separately for audio and audiovisual conditions, from a set of acoustic and phonetic features (sound envelope, phoneme onsets, and phonetic feature categories, see Figure 1B and C). Crucially, we considered neural activity to phonemic contrasts that are largely visually indistinguishable but auditorily distinct (i.e., *voicing*) in response to audio-only speech, and to contrasts that can be discriminated via visual cues (i.e., *place of articulation*) in response to audiovisual speech. Encoding of each additional feature was quantified as the gain in explained EEG activity obtained by stepwise addition of predictors to the model: first the gain provided by the phoneme onset compared to speech envelope, and subsequently the gain provided by auditory or visual phonetic features relative to the envelope + phoneme onset model.

Phoneme onset provides a general temporal cue, independent of specific phonetic features; therefore, it was expected to be robustly encoded across all groups, regardless of sensory experience during early development. In contrast, we predicted phonetic feature-specific effects: successful encoding of voicing during auditory speech was expected only in participants with access to speech sounds during the typical sensitive period for phonetic attunement (HC and AD), whereas successful encoding of place of articulation during audiovisual speech was expected across all groups, reflecting their access to visual phonetic features during the first year of life (see ^27,28^ for the encoding of visual speech features). These predictions test whether phonetic attunement reflects an intrinsically constrained yet modality-flexible developmental predisposition, rather than a process contingent on early auditory input.

## Results

We computed multivariate temporal response functions (mTRFs) at the single-participant level, separately for audio-only and audiovisual speech contexts, progressively adding predictors of interest to assess their specific contributions to model fit (Figure 1A and 2A; see ^3,29^ for a similar approach in adults and developmental populations). The predictors were entered incrementally into the models: first only envelope, then envelope + phoneme onset, then envelope + phoneme onset + phonetic features (auditory or visual). We measured EEG prediction correlations (Pearson’s *r*) for each model at the single-channel level; channel-wise tests were used to identify whether adding a predictor produced any measurable gain within a group and the resulting gain was averaged across significant channels. The effect of auditory experience rested on the planned comparisons between CI groups (AD vs. CD) and the feature-by-condition pattern (see in the Methods - Statistics - section for details). Comprehension accuracy was analyzed as an engagement check and is reported in the Supplementary Materials.

**Figure 2.**
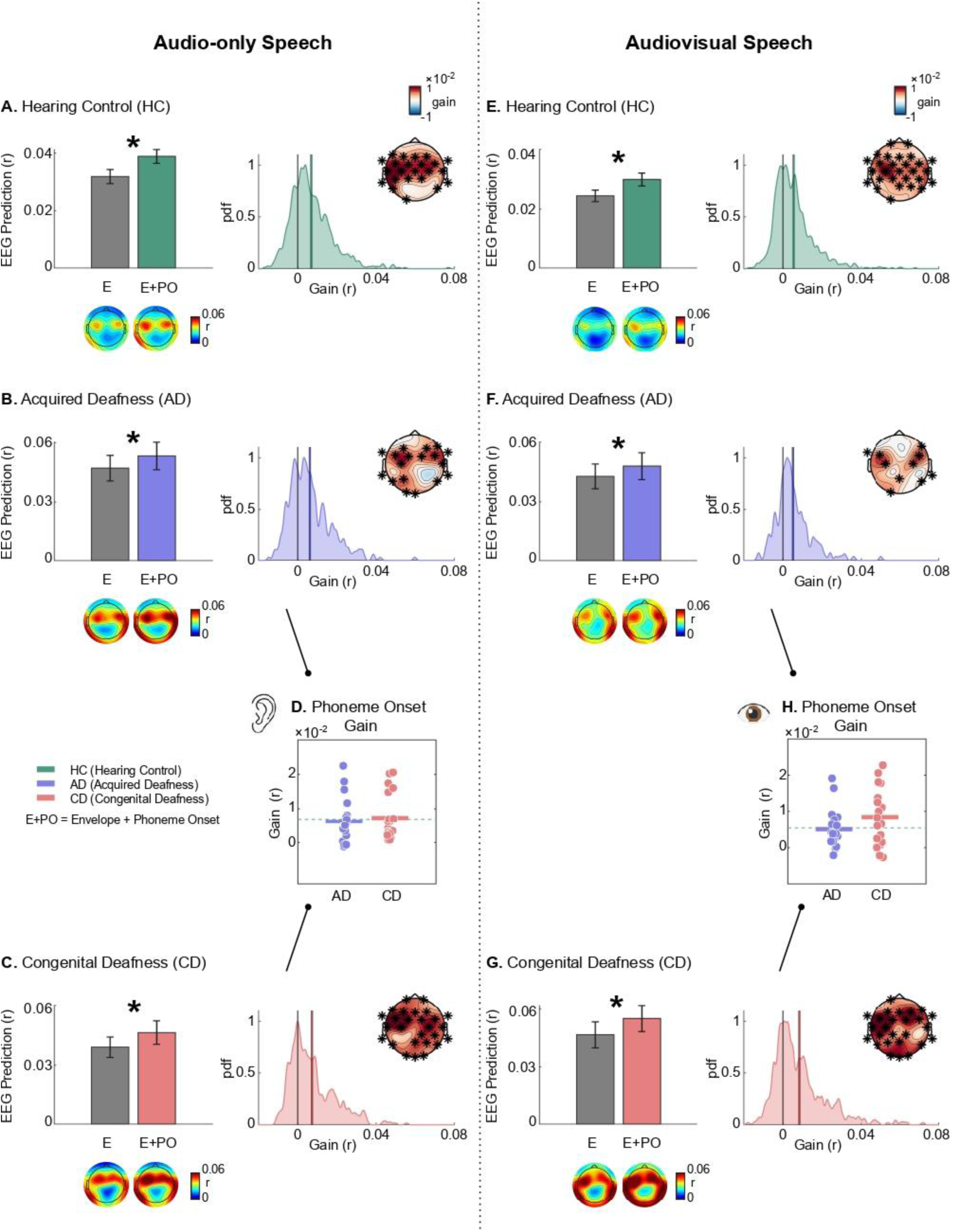
Neural Encoding of Phoneme onset. The left and right sides of the figure show results for the audio-only and audiovisual conditions, respectively. A) Phoneme onset gain for the HC group in the audio-only condition. Model prediction results are shown in the bar plot, for the Envelope-only (E) and the Envelope + Phoneme Onset (E+PO) models. The bar plot displays the group average of the mean prediction accuracy computed across significant channels at the individual level. Error bars represent the standard error. We marked the bar plot figure with an asterisk when at least one electrode showed a significant gain. Below, we show the topographies of the EEG prediction (r) for each model. The right panel shows the distribution of the pooled channel gains (pdf, probability density function); the topography displays the phoneme onset gain across the scalp, and asterisks highlight electrodes showing a significant gain. B) and C) Phoneme onset gain in the audio-only condition for the AD and CD groups, respectively. D) The plot shows the group averages of the phoneme onset gain measured in the AD and CD children, with dots representing individual participants. No significant difference emerged between the two CI groups. The green dashed line indicates the mean gain of the hearing control (HC) group. E), F) and G) Phoneme onset gain in the audiovisual condition for HC, AD and CD groups, respectively. H) The plot shows the mean gains in the AD and CD groups, with dots representing individual participants; no significant difference emerged. The green dashed line indicates the mean gain measured in the HC group.

### Phoneme onset

#### Audio-only

We examined the neural encoding of phoneme onsets during continuous speech processing within each group (i.e., HC, AD, and CD). In each group, adding the phoneme-onset predictor improved prediction accuracy at multiple electrodes relative to the envelope-only model (*speech envelope + phoneme onset > speech envelope-only;* one-tailed paired t-test at each electrode, α=0.05, uncorrected; see Figure 2A, B, and C). This pattern indicated robust encoding of a general temporal phonemic cue across developmental hearing histories. When the summary gain was compared between the two CI groups, CD and AD children did not differ (t(36)=-0.498, p=0.689, one-tailed; see Figure 2D). We therefore found no evidence that absence of auditory input during the first year reduced phoneme-onset encoding.

#### Audiovisual

Phoneme onset encoding during audiovisual speech mirrored the pattern observed in the audio-only condition. Adding phoneme onset improved prediction accuracy at multiple electrodes in all groups (Figure 2E, F, and G). The CD and AD groups did not differ in their summary gain (t(35)=-1.486, p=0.927, one-tailed; see Figure 2H) again indicating no detectable reduction in this general speech-timing response associated with early auditory deprivation.

### Phonetic features

#### Audio-only

CD children were not exposed to auditory speech during the sensitive period for the establishment of native phonetic categories. We therefore asked whether early auditory deprivation was associated with reduced encoding of a primarily acoustic phonetic feature: *voicing* contrasts between consonants matched for manner and place of articulation (see Methods). To this end, we tested for the existence of a *voicing* gain within each group (*speech envelope + phoneme onset + voicing* model > *speech envelope + phoneme onset* model). While HC and AD children showed a significant *voicing* gain at several electrodes (Figure 3A and B), no significant gain emerged in CD children (Figure 3C). No sensors showed a significant *voicing* gain in the CD group despite the permissive effect-detection threshold (uncorrected p-value used for within-group tests). To test the a priori CI contrast, we averaged voicing gain for both CI groups across the electrodes showing a voicing gain in the AD group and compared AD with CD. The *voicing* gain was greater in the AD group than in the CD group (*t*(36) = 2.492, p = 0.009, one-tailed; Cohen’s d = 0.78, 95th confidence interval = 0.22–1.36; see Figure 3D). Because both CI groups used implants at testing, this difference supports a selective association between auditory access during the first year of life and later audio-only encoding of auditory phonetic features.

**Figure 3.**
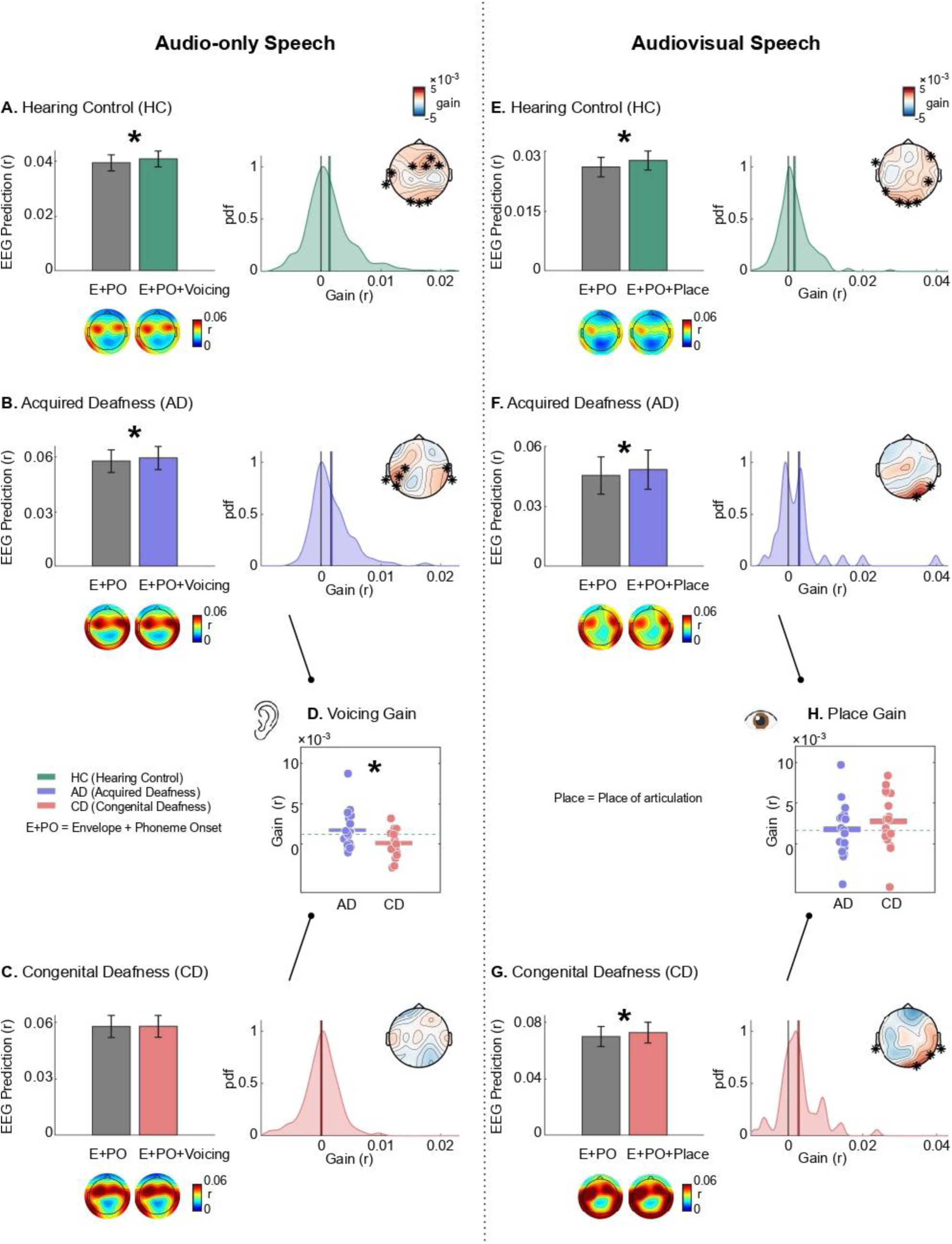
Neural Encoding of Phonetic features. The left and right sides of the figure show results for the audio-only and audiovisual conditions, respectively. A) Auditory phonetic feature (i.e., voicing) gain for the HC group in the audio-only condition. Model prediction results are shown in the bar plot, for the Envelope + Phoneme Onset (E+PO) and the Envelope + Phoneme Onset + Voicing (E+PO+Voicing) models. The bar plot displays the group average of the mean prediction accuracy computed across significant channels at the individual level; error bars represent the standard error. We marked the figure with an asterisk when at least one electrode showed a significant gain. Below, we show the topographies of the EEG prediction (r) for each model. The right panel shows the distribution of the pooled channel gains; the topography displays the auditory phonetic feature gain across the scalp, and asterisks highlight electrodes showing a significant gain. B) The figure shows the auditory phonetic feature gain for the AD in the audio-only condition. C) For the CD group, no significant auditory phonetic gain was observed at any electrode. D) The plot shows the group averages of the gain measured in the AD and CD children, with dots representing individual participants. The gain in the CD group was significantly lower than that in the AD group. The green dashed line indicates the mean gain measured in the hearing control (HC) group. E), F) and G) Visual phonetic feature (i.e., place of articulation) gain in the audiovisual condition for HC, AD, and CD groups, respectively. The bar plots show the model prediction results for the Envelope + Phoneme Onset (E+PO) and the Envelope + Phoneme Onset + Place of Articulation (E+PO+Place) models. H) The plot shows the mean gains in the AD and CD groups, with dots representing individual participants; the groups did not differ significantly. The green dashed line indicates the mean gain measured in the HC group.

#### Audiovisual

We next tested whether the reduced audio-only voicing gain in CD children reflected a general phonological-processing limitation or selective auditory phonetic features effect. To this end, we investigated the encoding of phonetic features that can be differentiated by visual cues (i.e., minimal pairs for *place of articulation*, see Methods section for more details on how the *place of articulation* predictor was created). CD children who could access visual speech during the sensitive period for the acquisition of phonetic categories might have attuned to such visual phonetic features. We tested for the existence of a *place of articulation* gain within each group (*speech envelope + phoneme onset + place* model > *speech envelope + phoneme onset* model). All groups showed a significant *place of articulation* gain (Figure 3E, F, and G), and no statistically significant differences were detected between the CD group and their AD peers (t(34)=-0.879, p=0.807, one-tailed; see Figure 3H). Results indicate that the CD group was able to successfully encode phonetic features that were provided by visual input during the sensitive period for phonetic attunement. Within CD children, the place-of-articulation gain in the audiovisual condition was larger than their audio-only voicing gain (t(17)=2.831, p=0.012). Thus, the CD group showed preserved encoding of phonetic structure when the relevant contrast was available through visual speech, arguing against a global inability to encode phonetic categories. Together, these findings support the conclusion that auditory deprivation during the first year of life selectively disrupts the neural encoding of acoustic phonetic features.

To assess feature and modality specificity, we also swapped predictors across conditions by testing *place-of-articulation* gain in audio-only speech and *voicing* gain in audiovisual speech. Testing whether the CD group failed to encode even *place of articulation* when it was conveyed exclusively by auditory cues would confirm their deficit in the processing of auditory phonetic features. Conversely, if *place of articulation* were successfully encoded by CD children during audio-only speech, this would indicate a crossmodal transfer in the acquisition of phonetic features from visual to auditory representations. The CD group did not show a significant gain for *place of articulation* when this feature was conveyed exclusively by auditory cues (see Supplementary Materials section - Place gain in Audio-only condition - section for further details and Figure S1). Results support the idea that discrimination between consonant contrasts strongly depends on the sensory phonetic features available during the sensitive period for phonetic attunement.

In addition, examining *voicing* gain in audiovisual speech allowed us to investigate whether CD children benefited from multisensory cues. A significant *voicing* gain emerged in all groups, with no difference observed as a function of auditory experience during the first year of life. This suggests a critical role of vision. Visual cues conveyed by coarticulation during continuous audiovisual speech processing may help CD children differentiate pairs of consonants that are not discriminable in audio-only speech (see Supplementary Materials section - Voicing gain in Audiovisual condition - section for further details).

## Discussion

Despite the biologically endowed capacity for speech acquisition, it remains unclear whether the sensitive period for phonetic attunement is triggered by the availability of auditory input. We addressed this question by examining the neural processing of phonetic features in children whose access to auditory speech had been restored through cochlear implants after a period of auditory deprivation, as well as in hearing controls. We specifically contrasted the neural activity of CI children who experienced auditory deprivation during the first year of life (i.e., CD) with that of their CI peers who received auditory input during this phase (i.e., AD). Neural responses were recorded during continuous auditory and audiovisual speech processing, enabling the assessment of encoding of both auditory and visual phonetic features. Results showed that only children who had access to hearing during the sensitive period for phonetic attunement (i.e., HC and AD) successfully encoded auditory phonetic features, whereas children who were auditorily deprived during this time window (i.e., CD) did not. However, CD children benefited from the access to visual phonetic features, similarly to their AD and HC peers. These data indicate that the sensitive period for phonetic attunement is not triggered or fully determined by the presence of auditory input; rather, the underpinning neural circuitry exploits the available speech cues, which, in the case of children with congenital deafness, are conveyed through sight. These findings are consistent with the view that postnatal auditory experience refines, rather than creates, speech-processing pathways ^30^.

Our data support the hypothesis that the sensitive period for phonetic attunement is not contingent on auditory experience alone; once this developmental time window has passed, restoring auditory input in the case of deprivation no longer leads to full tuning to speech sounds. Previous evidence showed that early or increased auditory experience does not trigger an earlier opening of this sensitive period in preterm infants ^14^, supporting the idea that the timing of speech perception development is tightly constrained by brain maturation. However, animal studies have shown that sensory deprivation can delay or pause the development of perceptual functions until crucial sensory stimulation is provided (e.g., ^31,32^). Our data reveal that, when auditory deprivation extends across at least the first year of life, the absence of auditory stimulation alone is insufficient to maintain open critical periods for the perceptual foundations of language, such as phonetic attunement. Rather, severe reductions in social and linguistic input across modalities may more profoundly delay language-related critical periods, as observed for phonetic attunement in infants of untreated depressed mothers ^13^ or in the case of infants suffering social deprivation ^33,34^. Consistent with this view, animal studies have shown that critical periods for communication-related functions, such as birdsong learning, can remain open if the animal is reared in isolation ^35^.

Our results demonstrate that early auditory deprivation differentially shapes the neural processing of auditory and visual phonetic features. In the case of congenital deafness (CD), consonant contrasts that are differentiable primarily via their acoustic phonetic features are especially challenging for CI children to encode. This context is analogous to what occurs with non-native sound contrasts in hearing children: native English speakers fail to discriminate Hindi dental /d̪a/ versus retroflex /ɖa/ given their lack of exposure to these speech statistics during the sensitive period for phonemic attunement ^5^. In contrast, CI children who had access to auditory speech patterns during the first year of life and acquired deafness later in development (the AD group) were able to successfully exploit acoustic phonetic differences. Their successful neural processing of these features confirms that auditory stimulation received through CIs is sufficient to detect phoneme differences, and that the absence of such neural encoding in the CD group is rather the effect to the lack of experience with sounds in early development. Overall, these findings align with studies of international adoptees showing the strong impact of early experience. Even years later, as adults, adoptees retain sensitivity to phonetic patterns they perceived during roughly the first year of life, despite having been adopted into an environment where they were no longer exposed to those phonemes ^36^.

The data also revealed that impairment in neural processing in infants with congenital deafness is selective to auditory phonetic contrasts and does not reflect a general phonological processing deficit. All groups benefitted from visual phonetic information during audiovisual speech processing, indicating that for visually distinguishable consonant pairs, such as /m/ and /n/, visual input during early life provides sufficient speech information to detect a bimodal distribution, which in turn also allows CD infants to establish separate phoneme categories. Although phonology is traditionally considered to be primarily sound-dependent, visual phonetic information is also discriminated very early in development ^17^. Consistent with the intrinsically multimodal nature of language, the sensitive period for phonetic attunement in auditory-deprived infants was shaped by visual speech statistics when vision was the only available modality. However, the data suggest a modality-specific acquisition of visual and auditory phoneme categories. In fact, the CD children did not show successful neural encoding of phonemes that are visually distinguishable (e.g., /m/ and /n/) when provided only acoustic signals (see supplementary Materials - Place of articulation in the Audio-only condition - section). This points to an ineffective transfer or interplay of phonetic representations across sensory channels. Moreover, it is important to note that CD infants, like the other groups, showed successful neural encoding of phoneme onsets not only when visual information was available but also during auditory-only speech exposure. This may reflect that phoneme-onset encoding relies on simpler processing mechanisms compared with the processing of linguistically meaningful phonetic features, or that auditory and visual cues characterizing phoneme onset are highly correlated ^37^. Perhaps, CD children can take advantage of speech cues that are coherent across sensory modalities, when audiovisual consistency is high (see also Supplementary Materials - Voicing gain in the Audiovisual condition - section).

Given the heterogeneous speech processing performance of children with cochlear implants ^38^, shedding light on the impact of auditory deprivation timing on different aspects of natural speech processing is fundamental for improving and tailoring interventions. Whereas basic speech-processing mechanisms do not appear to be specifically disrupted by auditory deprivation during the first year of life, as previously shown for speech-envelope tracking ^39^ and here for the encoding of phoneme onsets, the processing of specific phonetic features is clearly shaped by early experience. This vulnerability in phonetic processing aligns with evidence of phonological impairments in children with cochlear implants ^40,41^. Such a deficit may contribute to downstream effects on language development, as the perception of speech features plays a central role in the acquisition of more complex linguistic abilities ^42^, in particular, neural responses to consonants have been shown to predict lexical acquisition ^43^. Given the importance of the first months of life for the formation of meaningful categories as a function of sensory experience and the severe consequences of early auditory deprivation ^44^, future studies should investigate how to enhance tuning to these highly vulnerable phonetic features. This is necessary to support establishing the full native phoneme inventory in infants with congenital deafness and ensuring long-lasting effects on neural processing of continuous speech.

### Limitations

Our results clearly indicate that when there is auditory deprivation, the sensitive period for phonetic attunement is not simply paused or held open until auditory input is introduced, highlighting the importance of providing access to auditory speech signals during the first year of life. Yet, it remains possible that more subtle forms of flexibility are retained beyond infancy ^45^. However, longitudinal studies are required to determine the extent to which plasticity supporting the development of auditory phonetic tuning persists. Testing CI individuals at different time points following implantation would help clarify the extent to which auditory phonetic tuning is possible, with important clinical implications.

A second limitation of this study is the absence of a visual-only condition. Without a unimodal visual speech condition, we cannot fully disentangle whether CD children’s ability to encode visual phonetic features during audiovisual speech reflects the mere presence of unisensory visual cues or a benefit due to multisensory integration. Nevertheless, the present findings demonstrate that visual phonetic information available during the first year of life is sufficient to support the formation and long-term neural encoding of visually discriminable phoneme categories, even in the absence of early auditory input. To promote speech processing in CI children born with congenital deafness, further studies are required to clarify the underlying mechanisms and determine whether and how early auditory deprivation alters multisensory pathways ^46^.

### Conclusion

These findings reveal that when infants experience auditory deprivation during the sensitive period for native phonetic categories, perceptual attunement is not postponed until access to auditory input is restored. Instead, attunement progresses by relying on the phonetic information available during early development. Accordingly, in the context of congenital deafness, infants tune in to phonetic features conveyed through visual cues. Thus, phonetic attunement does not depend on auditory experience per se, but it does depend critically on early access to informative sensory evidence.

## Methods

### Participants

A total of 93 children participated in the study. Fifty-six children with cochlear implants were recruited at the Meyer Hospital of Florence (Italy) and the IRCCS Materno Infantile Burlo Garofolo of Trieste (Italy). In line with our main goal, we collected the following clinical variables to characterize CI participants: results of the newborn hearing screening with otoacoustic emissions (pass or fail), hearing thresholds in each ear to determine the level of deafness (moderate, severe, or profound), and age at cochlear implantation. This information allowed us to identify a group of congenitally deaf CI children who were auditory deprived during the first year of life, and a group of CI children with acquired deafness who were exposed to auditory environmental statistics during the first year of life (See Supplementary Materials “Criteria for CD and AD groups classification”). The selected CI individuals with congenital bilateral deafness (hereafter CD) received a cochlear implant at the earliest at 11 months of age, whereas all CI individuals with acquired deafness (hereafter AD) received a diagnosis of profound bilateral deafness after twelve months of age (See Supplementary Materials “Participant excluded” for details on participants were not included in the analysis). We analysed nineteen CD children (age range: 2.92-14.58 years, mean age=8.99; SD=3.50, nine females and ten males) and nineteen AD children (age range: 4.50-16.33 years, mean age=8.54; SD=3.20, eleven females and eight males). Consistent with group selection criteria, CD children received a diagnosis of profound bilateral deafness significantly earlier than their AD peers (t(18.02)=-7.906, p<0.001). Consequently, the CD group had significantly more experience with the implant compared to the AD group (t(36)=2.158; p=0.038, CD mean: 78.84 months of age SD=36.75, AD=51.53, SD= 41.15). The median age at cochlear implantation was 15 months (IQR=11, range 11-132) for the CD group and 42 months (IQR=32, range: 17-120) for the AD group. Importantly, all CI users received their cochlear implant at least six months before the experimental session to ensure a consolidated auditory experience ^47^. Note that both participants with bilateral and unilateral CI are included (CD thirteen bilateral, AD nine bilateral). Additionally, hearing children (HC, N=37; range: 3.50-18.75 years, mean age=9.04 years; SD=4.10, seventeen females and twenty males) were recruited as an age- and sex-matched control group with typical auditory experience during development. There are no significant differences across the three groups in age (F(2,72)=0.117; p=0.890) or sex distribution (χ2(2)=0.756; p=0.685). HC children were recruited at IMT School for Advanced Studies in Lucca (Italy).

Classification into CD and AD groups was performed before EEG analysis using exclusively clinical history. CD status required failed neonatal otoacoustic-emission screening and objective diagnosis of profound bilateral deafness within two months of birth. AD status required evidence of hearing or non-profound hearing loss during the first year and diagnosis of profound bilateral deafness after 12 months of age. CI children with ambiguous auditory history were excluded before analysis (see Supplementary Materials).

None of the children included in the final sample had additional sensory deficits or neurological disorders, as confirmed by medical records and/or family reports. All CI participants were trained to oral and not sign language. For all participants Italian was first language (L1), except for six children who were bilingual (one CD, two AD, and three HC participants). The study was approved by the local Ethics Committees (“Comitato Etico congiunto per la ricerca della Scuola Normale Superiore e della Scuola Superiore Sant’Anna” approval number 34/2020, and “Comitato Etico Regione Toscana - Pediatrico dell’Azienda Ospedaliera Universitaria Meyer” approval number 17/2020). Before participation, written informed consent was obtained from parents and from children older than seven years of age. The experimental protocol adhered to the principles of the Declaration of Helsinki (2013), and all ethical regulations relevant to human research participants were followed.

### Stimuli

The speech stimuli consisted of 3-minute stories read by a native Italian speaker with formal diction training. To ensure age-appropriate content, we selected different stories based on the children’s age groups, corresponding to Italian school stages (3–6, 7–10, and 11–15 years). For each group, ten stories were chosen from Italian children’s books suitable for that developmental stage. All recordings were made in a sound-attenuated booth (BOXY, B-Beng s.r.l., Italy) using an iPhone 7 (12 MP camera, video resolution in HD, 720p with 30 fps, at a sampling frequency of 48 kHz) connected to an external condenser microphone (YC-LM10 II, Yichuang). The video comprised the speaker’s face recorded with a homogeneous grey wall in the background. We created two versions of each story: an audio-only and an audiovisual version. The stories were balanced across conditions within each subgroup. After brief preprocessing (see Supplementary Materials of ^39^), the stories were presented to participants using PsychoPy® software (PsychoPy3, v2020.1.3). Speech audio was delivered through a single front-facing loudspeaker (Bose Companion® Series III, USA) positioned approximately 70 cm from the participant’s head, behind the computer screen on which the video was displayed. Participants sat approximately 60 cm from the screen. The volume was calibrated to approximately 80 dB SPL using a Meterk MK09 Sound Level Meter.

### Task and experimental procedure

Participants were instructed to listen attentively to the stories while facing a computer screen positioned in front of them. At the beginning of each story, a white fixation cross appeared on the screen. After two seconds of silence, the title of the story was displayed, followed by the start of the audio narration. In the audio-only condition, the fixation cross remained at the center of the screen throughout the story and changed color at random intervals (every 1 to 20 seconds) to help maintain the children’s gaze fixation on the cross, whereas in the audiovisual condition, the speaker’s face was displayed on the screen. Throughout the recording, an experimenter was present to ensured that children were attending to the screen. At the end of each story, participants completed a brief comprehension task consisting of two-alternative forced-choice (2AFC) questions designed to assess their understanding and encourage active listening (see ^39^ for further details on the questions and Supplementary Materials for the mean groups data within each condition). The order of conditions was fixed: first the audio-only condition and then the audiovisual condition. This was chosen to begin with the less engaging condition and finish with the more engaging one, when children might be more tired. The fixed order was chosen before data analysis. It makes the audiovisual condition potentially more susceptible to order or fatigue effects, but it cannot explain the critical audio-only voicing result because the audio-only condition was always administered first. For each condition, participant listened to four different stories. However, due to limited participation, eleven hearing control (HC) children and ten cochlear implant (CI) users listened to only three stories in the audio-only condition (4 CD and 6 AD); in the audiovisual condition, eight HC and thirteen CI participants completed three stories (5 CD and 8 AD). Electroencephalographic (EEG) activity was recorded continuously throughout the entire experimental session. Note that one HC participant and one CD participant ended the experiment after the audio-only condition, missing the audiovisual condition.

### EEG recording and preprocessing

EEG data were recorded using a Brain Products ActiCHamp Plus system with 32 active electrodes mounted on elastic caps (EasyCap Standard 32Ch actiCAP snap), specifically adapted for children. Signals were sampled at 500 Hz and recorded using Brain Vision Recorder software. For participants with cochlear implants (CI), electrodes located in close proximity to the implant magnet were disconnected (mean number of disconnected electrodes = 3.50, SD = 1.44; range: 1–7). EEG data from each story presentation were concatenated and preprocessed offline using the EEGLAB toolbox (version 14.1.2) in MATLAB 2019b, following a validated artifact removal pipeline. The continuous EEG data were first low-pass filtered at 40 Hz (Hanning window, filter order = 50), downsampled to 250 Hz, and then high-pass filtered at 1 Hz (Hanning window, filter order = 500). The resulting signal was segmented into consecutive 1-second epochs, and noisy segments were automatically rejected based on joint probability (threshold set at 3 standard deviations across all channels ^48^). To remove prototypical artifacts such as eye blinks and eye movements, Independent Component Analysis (ICA) using the extended Infomax algorithm was performed ^49–51^. The ICA decomposition was computed on the filtered and downsampled data without noisy segments to improve signal processing ^52^. Resulting ICA weights were applied to the continuous raw original data to retain signal fidelity ^53^. Artifact components were identified and removed using CORRMAP (version 1.03 ^54^), a semi-automated procedure in which prototypical topographies for blink and eye movement artifacts were selected as templates. Components showing a correlation greater than 80% with the template were excluded from the data. The mean number of removed components was 2.00 (SD = 0.00) for the HC group and 2.09 (SD = 0.39) for the CI group in the audio-only condition, and in the audiovisual condition, 1.94 (SD = 0.34) SD for HC and 1.86 (SD = 0.48) for CI participants. Moreover, since EEG recordings from CI users are affected by electrical artifacts generated by the implant, these artifacts were removed with a method suitable for neural speech tracking analyses. Given that CI artifacts are expected to peak around 0 ms lag, we used a combination of Second Order Blind Identification (SOBI) and a classification algorithm to isolate and remove artifact-dominated components. Specifically, we: (i) decomposed the EEG using SOBI to separate signal and noise sources; (ii) applied the Temporal Response Function (TRF) model to each SOBI component to obtain SOBI-component-TRFs; (iii) modeled these TRFs with Gaussian fits (one peaking before 0 ms, up to five after); and (iv) identified artifact components based on parameters (i.e., the R² ratio and beta value of the pre-zero Gaussian) from the HC group using a decision boundary that limited false positives to 5%. Identified artifact components were then excluded, and the cleaned EEG signal was reconstructed (see Federici et al., 2025 for more details on this method). In audio-only condition we removed on average 2.69 (SD = 2.39) for the CD group, and 2.25 (SD = 2.02) for the AD group, and in audiovisual condition 1.45 (SD = 0.52) for CD and 1.92 (SD = 1.25) for AD.

After removing ICA components (for prototypical artifacts) and SOBI components (for CI-related artifacts), the artifact-cleaned, unfiltered EEG data were processed as follows: low-pass filtered at 40 Hz (Hanning window, order 50), downsampled to 250 Hz, and high-pass filtered at 0.1 Hz (Hanning window, order 5000; see Federici et al., 2025). Noisy channels were detected using the clean_channels function from the clean_rawdata EEGLAB plugin (correlation threshold = 0.8; sample size = 1; other parameters at default) and interpolated via spherical spline interpolation (mean ± SD interpolated channels: in audio-only condition HC = 2.30 ± 1.15; CI = 1.88 ± 1.60; and in audiovisual condition HC = 2.05 ± 1.64, CI = 2 ± 1.41). Electrodes disconnected near the CI magnet were also interpolated. The data were then re-referenced to the average reference. We further band-pass filtered according to the frequence of interested, between 1–8 Hz (high-pass: 1 Hz, order 500; low-pass: 8 Hz, order 126; Hanning windows), consistent with prior work demonstrating that phoneme encoding is primarily reflected in low-frequency neural activity ^3,29^. Within each condition, the EEG from each story was epoched into 2.5-minute segments (excluding initial seconds to avoid onset responses; see Supplementary Materials 1.3.2), downsampled to 100 Hz, and divided into 50-second trials. This yielded 12 trials per participant (or 9 for those who listened to only three stories) for each condition. Trials were used for cross-validation, and data were z-scored to optimize regularization during model fitting ^55^. All preprocessing and artifact-removal steps were applied before fitting phonetic-feature models and were identical for HC, AD, and CD children, except for the CI-artifact correction that was applied only to CI users. The CI-artifact classifier was defined using the HC group to limit false positives, reducing the risk that group differences in phonetic-feature gains were introduced by differential artifact removal. Disconnected and noisy channels were interpolated before average referencing; because summary gains were based on cross-validated prediction accuracy, residual artifacts would have had to selectively mimic phonetic-feature timing to account for the reported effects.

### Extraction of the speech features

For each story, the acoustic envelope was computed by taking the absolute value of the Hilbert transform and applying a low-pass Butterworth filter (3rd order, 8 Hz cut-off; MATLAB filtfilt function). Phoneme onsets and their articulatory features—manner and place of articulation, and voicing—were extracted using Praat and manually verified for timing accuracy using MAUS software ^56^. A phoneme onset predictor was created as a pulse vector, with ones at the latency of each phoneme onset and zeros at the remaining samples. To construct the voicing predictor, i.e. a selectively acoustic feature, we selected consonant pairs that differed only in voicing while sharing the same manner and place of articulation. These voicing minimal pairs were /d/–/t/, /b/–/p/, /g/–/k/, /z/–/s/, /dz/–/ts/, and /v/–/f/.

We then generated a two-column matrix of pulse vectors: the first column contained ones at the onset of voiced consonants, and the second column at the onset of voiceless consonants. To construct the place of articulation predictor, we selected consonant pairs that differed only in place of articulation while sharing the same manner of articulation and voicing. These contrasts were chosen because the relevant articulatory distinction (alveolar versus labial/labiodental constriction) is visually accessible during face-to-face speech. The predictor therefore indexes a phonetic category whose realization can be supported by visual speech in the audiovisual condition. The selected place contrasts were /n/–/m/, /s/–/f/, /z/–/v/, /d/–/b/, and /t/–/p/. We then generated a two-column matrix of pulse vectors: the first column contained ones at the onset of alveolar consonants, and the second column at the onset of labial or labiodental consonants. For each participant, speech features from the stories were concatenated in the order of presentation and segmented into 50-second trials, yielding twelve trials per participant (or nine for those who listened to only three stories). All features were downsampled to 100 Hz to match the EEG data. The envelope was normalized by rescaling its values between 0 and 1 to ensure that predictors had comparable magnitudes.

### The forward models

To investigate how children’s brains encode speech features within each condition (i.e., audio-only and audiovisual), we employed a linear forward model known as temporal response function (TRF), implemented using the mTRF toolbox (Version 2.4 ^55^). Separate TRF models were fitted at the individual subject level to predict the neural response at each of the 32 EEG channels from the acoustic features of interest, using a time-lag window ranging from −100 to 600 ms in 10 ms steps. Three encoding models with increasing levels of speech-feature complexity were implemented to assess predictions performance in stepwise fashion. The first-level model, serving as a baseline model, included only the speech envelope as a predictor. The second-level model included the speech envelope and phoneme onset (hereafter, *speech envelope + phoneme onset* model). Finally, the third-level model included either the voicing predictor in addition to the envelope and phoneme onset (hereafter, *speech envelope + phoneme onset + voicing* model), or the place of articulation predictor in addition to the envelope and phoneme onset (hereafter, *speech envelope + phoneme onset + place* model). Each model was trained using a leave-one-out cross-validation procedure: all trials except one were used to train the model to predict neural responses from the given speech features (i.e., envelope; envelope and phoneme onset; envelope, phoneme onset, and voicing or envelope, phoneme onset and place of articulation), and the remaining trial was used for testing. This process was repeated for all trials, and the resulting models and their prediction accuracies (r) were averaged within each EEG channel. Importantly, the regularization parameter (λ) was estimated for each training fold by selecting the λ value that minimized the mean squared error (MSE).

### Statistics

To assess whether the inclusion of each additional speech feature significantly improved EEG prediction accuracy (i.e., the correlation coefficient *r* between predicted and actual EEG signals), we compared each model with its previous version, with a stepwise approach. First, we assessed the phoneme onset gain during audio-only and audiovisual speech processing by contrasting the *speech envelope + phoneme onset* model accuracy with the *speech envelope* model accuracy, separately for audio-only and audiovisual conditions. Next, we assessed auditory and visual phonetic features gain during audio-only and audiovisual speech processing, respectively. The *speech envelope + phoneme onset + voicing* model accuracy was contrasted with the *speech envelope + phoneme onset* model accuracy in the audio-only condition to assess the gain induced by the auditory phonetic features, and the *speech envelope + phoneme onset + place* model accuracy was contrasted with the *speech envelope + phoneme onset* model accuracy in the audiovisual condition to assess the gain induced by the visual phonetic features. Finally, we also investigated the place of articulation gain in the audio-only condition and the voicing gain in the audiovisual condition (see Supplementary “Place of articulation in the Audio-only condition” and “Voicing gain in the Audiovisual condition”).

To assess the existence of each gain within each group (HC, AD, and CD), we tested at each electrode whether the inclusion of a specific speech feature significantly improved prediction accuracy. After applying Fisher’s z-transformation to the *r* values to approximate a normal distribution, one-tailed paired t-tests (α = 0.05) were performed, because all nested-model comparisons had a single meaningful direction: an informative feature should improve cross-validated prediction relative to the corresponding model without that feature. These electrode-wise tests were used as descriptive effect-detection maps and were not interpreted as familywise-corrected claims about scalp topography. Because we had no a priori hypothesis regarding which channels would show a gain, and because spatial effects could vary across groups, we selected channels showing a significant gain at the group level to compute the average gain at the individual level. This permissive threshold was chosen to detect even small gains within each group. The critical developmental inference was the planned comparison between CD and AD children, because both groups were CI users at test and differed in auditory access during the first year of life. One-tailed independent-samples t-tests were used for these planned comparisons with the directional alternative CD < AD. This direct comparison between the two CI groups allowed us to control for factors related to cochlear implantation per se and to attribute any reduction in gain specifically to differences in auditory experience during the first year of life, which is what distinguishes the AD and CD groups. For the audio-only voicing test, the mask was defined from electrodes showing a positive voicing gain in the AD group, which served as the CI positive-control group with early auditory experience; this analysis asked whether CD children expressed the same response in sensors in which CI users with early auditory access encoded the feature. Before conducting group comparisons, we screened for within group outliers for each measured gain (mean ± 3 SD). In case outliers were detected, analyses were run excluding those values. However, analyses including outliers were also reported in the Supplementary Materials to show that the results were not driven by their presence or absence (see Supplementary Materials - Analysis including the outliers - section).

Model comparisons were nested and cross-validated: a feature was considered encoded only when adding it improved prediction. The same model architecture, lag window, regularization procedure, and cross-validation scheme were applied to HC, AD and CD participants within each comparison, so group differences in gain cannot be attributed to different model complexity.

## Acknowledgements

We thank all the children and their families who participated in this study; Erica Iob for assisting with data collection for some participants; Maria Aisa for organizing clinical information; and Sara Pintonello and Costanza Fratini for patient recruitment.

Funding: Ministero dell’Istruzione, dell’Università e della Ricerca (MIUR), PRIN 20177894ZH to D.B.; grant of Intesa Sanpaolo Bank to D.B.; Cochlear Technology Centre Belgium co-funded F.C.

## Authors contributions

Conceptualization: A.F., D.B.

Methodology: A.F., D.B., G.D.L., G.H., C.B., M.F.

Investigation: A.F., M.F., F.C.

Visualization: A.F., D.B, G.H.

Funding Acquisition: D.B.

Project Administration: D.B.

Supervision: D.B.

Writing–Original Draft: A.F., D.B.

Writing–Review and Editing: A.F., D.B., G.D.L, G.H., L.P., J.W., M.F., F.C., C.B., E.O., B.B

## Data availability

De-identified processed EEG data and speech-feature matrices required to reproduce the reported analyses will be deposited in an open repository before submission. Raw EEG data and individual clinical histories may require controlled access because the sample includes paediatric cochlear-implant users with rare clinical histories. Repository accession numbers and any access conditions should be added before submission.

## Code availability

Analysis scripts implementing mTRF modelling and statistical tests will be deposited in the same repository before submission. Custom code should be documented sufficiently to reproduce all reported figures and statistics.

