## Supplementary Materials for "Visual speech supports phonetic attunement in the absence of early auditory input"

**Place of articulation in the Audio-only condition**

We investigated the existence of *place of articulation* gain within each group also in audio-only speech. Since no visual stimulus was presented to participants, here we assessed auditory phonetic features of place of articulation (e.g. /n/ vs. /m/; alveolar vs. labial pairs). Within each group, we tested at each electrode whether the prediction accuracy of the *speech envelope + phoneme onset + place of articulation* model was significantly higher than that of the *speech envelope + phoneme onset* model. Only in the HC and AD groups did a significant *place of articulation* gain emerge (see Figure S1 panels A and B, note that place of articulation is more difficult, and for HC group only a tendency emerged $p<0.07$), while no gain emerged in the CD group (Figure S1 panel C). We performed a t-test to assess whether the AD group showed a higher gain with respect to the CD group. Results revealed that the CD children had a significantly lower gain with respect to their AD peers ($t(36)=4.762$, $p<0.001$, one-tailed, Cohen's $d = 1.51$, 95th confidence interval = 0.80–2.29; see Figure S1 panel D). These results confirmed that children who were auditorily deprived during the sensitive period for acquisition of native phonetic categories are not able to encode auditory phonetic features that can instead be encoded when visual cues are present (results of place of articulation in audiovisual speech).

**Voicing gain in the Audiovisual condition**

Finally, we explored voicing gain within each group also when participants processed audiovisual speech. We tested at each electrode whether the prediction accuracy of the *speech envelope + phoneme onset + voicing* model was significantly higher than that of the *speech envelope + phoneme onset* model. As expected, in both HC and AD groups a significant voicing gain emerged (see Figure S1 panels E and F). However, surprisingly, a gain also emerged in the CD group, and no statistically significant differences were detected between the CD and AD group ($t(33)=0.114$, $p=0.455$, one-tailed; see Figure S1 panels G and H). These results suggested that for CD children, exposure to visual information could help identify subtle differences between voicing minimal pairs of consonants. We speculate that this may be due to coarticulation during continuous speech processing, which might uncover visual differences between voicing minimal pairs. This result highlighted the importance of multisensory information during naturalistic speech processing. Moreover, some voicing cues (e.g., VOT) are subtle temporal cues that can be picked up visually. Finally, it should be acknowledged that the minimal residual hearing that is often available to deaf listeners before implantation could be enough to integrate with these visual cues and might have played a role. This highlights the importance of optimizing the residual hearing of these children while they are waiting to get an implant.

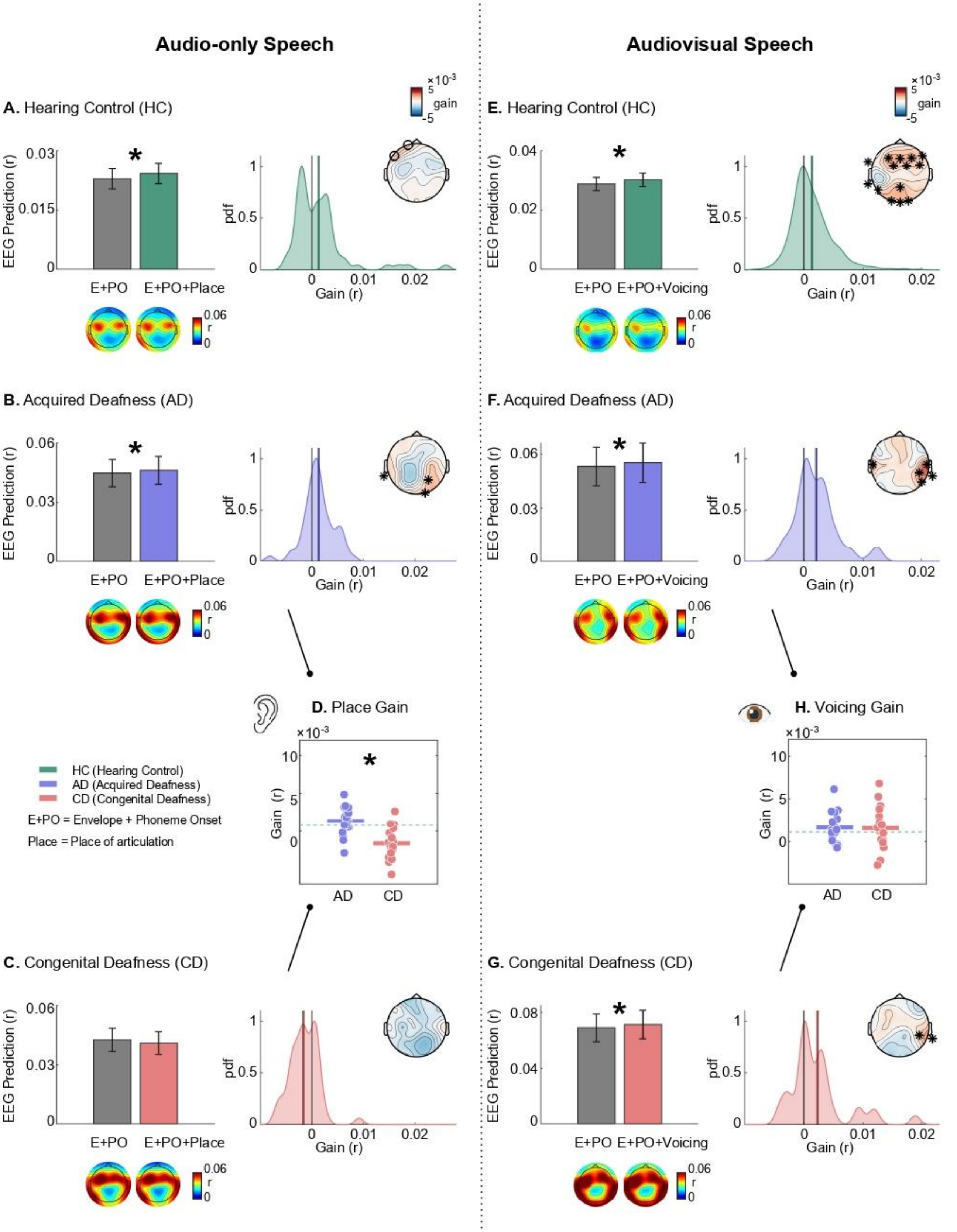


Figure S1. The left and right sides of the figure show results for the place of articulation gain in audio-only and the voicing in audiovisual conditions, respectively. A) Place of articulation gain in audio-only condition for the HC group in the audio-only condition. Model prediction results are shown in the bar plots, for the Envelope + Phoneme Onset (E+PO) and the Envelope + Phoneme Onset + Place of articulation (E+PO+Place)

models. Bar plot displays the group average of the mean prediction accuracy computed across significant channels at the individual level. Error bars represent the standard error. We marked the figure with an asterisk when at least one electrode showed a significant gain. Below, we show the topographies of the EEG prediction (r) for each model. The right panel shows the distribution of the pooled channel gains; the topography displays the gain across the scalp, and circles highlight electrodes showing a tendency toward a significant gain. B) The figure shows the place of articulation gain for the AD in the audio-only condition; the topography displays the gain across the scalp, and asterisks highlight electrodes showing a significant gain. C) No place of articulation gain was observed for the CD in the audio-only condition. D) The plot shows the group averages of the gain measured in the AD and CD children, with dots representing individual participants. The gain in the CD group was significantly lower than in the AD group. The green dashed line indicates the mean gain of the hearing control (HC) group. E), F) and G) Voicing gain in the audiovisual condition for HC, AD and CD groups, respectively. The bar plots show the model prediction results for the Envelope + Phoneme Onset (E+PO) and Envelope + Phoneme Onset + Voicing (E+PO+Voicing) models. H) The plot shows the mean gains in the AD and CD groups, with dots representing individual participants; the groups did not differ significantly. The green dashed line indicates the mean gain measured in the hearing control (HC) group.

### Analysis including the outliers

#### Phonetic features gain

**Place gain in Audiovisual condition.** One outlier emerged in the AD group. Including this participant did not change the result ($t(35)=0.088$, $p=0.465$).

**Voicing gain in Audiovisual condition.** One outlier was identified in the AD group and one in the CD group. Including both participants did not change the result ($t(35)=-0.161$, $p=0.563$).

### Criteria for CD and AD groups classification

Congenital deafness (CD) was ensured by the following criteria: (a) having failed to pass the neonatal screening for otoacoustic emissions (typically < 1 week after birth); (b) receiving a diagnosis of profound bilateral deafness (hearing thresholds ≥ 90 dB in both ears) following the objective evaluation of auditory brain-stem responses (ABR) within two months of age. Both these criteria are needed to classify a participant as a CD. Moreover, only CD children implanted after 11 months of age were selected to ensure auditory deprivation during the sensitive period for phonetic attunement. In contrast, AD children had at least some auditory experiences in early development (i.e., a minimum of 12 months). To ensure the presence of auditory experience during the first year of life, we combined the following clinical information: (a) whether they passed otoacoustic emissions neonatal screening at least with one ear; (b) an ABR indicating normal hearing before the diagnosis of deafness or a diagnosis of deafness that was not profound bilaterally (e.g., moderate deafness at least in one ear) made by ABR or behavioral test (hearing thresholds < 90 dB in at least one ear); (c) family report indicating residual hearing for the first period of life.

**Participant excluded**

Six CI participants were excluded for several reasons: (i) one two-year-old child was removed because they were unable to complete the experimental session; (ii) two children were excluded due to IQ scores falling below the age-normative range; (iii) one child underwent reimplantation several years after the initial implantation as a consequence of an infection; (iv) one child was excluded because of poor EEG data quality (15 electrodes were identified as bad channels); and (v) one participant was excluded due to technical issues during data acquisition.

The remaining fifty CI participants were categorized based on the timing of deafness onset and thus their auditory experience during the first year of life. We classified as CD children with profound bilateral congenital deafness who had their auditory function restored at the end of the first year of life (cochlear implant ≥ 11 months of age) and AD children who developed profound bilateral deafness after the first year of life (> 11 months of age). Twelve CI children were not included in the final analyses because auditory history did not allow clear classification into either group. Two participants received the implant before 11 months of age; one child acquired profound bilateral deafness at 7 months; and for nine children the onset of profound bilateral deafness was unclear.

**Behavioral data**

We computed the percentage accuracy on the story comprehension test for each participant separately in audio-only and audiovisual conditions. These data were used as an engagement check and are reported descriptively because the neural hypotheses concerned model-based encoding of speech features rather than explicit comprehension. In the audio-only condition, the HC group had a mean accuracy of 85.42 (SD = 15.22), the AD group 63.16 (SD = 17.53), and the CD group 58.99 (SD = 22.06). In the audiovisual condition, the HC group had a mean accuracy of 86.23 (SD = 15.53), the AD group 75.00 (SD = 20.81), and the CD group 78.24 (SD = 17.18).
